# Rapamycin attenuated zinc-induced tau phosphorylation and oxidative stress in animal model: Involvement of dual mTOR/p70S6K and Nrf2/HO-1 pathways

**DOI:** 10.1101/2021.09.24.461637

**Authors:** Chencen Lai, Yuanting Ding, Qian Chen, Songbai Su, Heng Liu, Ruiqing Ni, Zhi Tang

## Abstract

Alzheimer’s disease is pathologically featured by abnormal accumulation of amyloid-beta plaque, neurofibrillary tangles, oxidative stress, neuroinflammation, and neurodegeneration. Metal dysregulation including excessive zinc released by presynaptic neurons plays an important role in tau pathology and oxidase activation. The activities of mammalian target of rapamycin (mTOR)/ ribosomal S6 protein kinase (p70S6K) are elevated in the brains of patients with Alzheimer’s disease. Zinc induces tau hyperphosphorylation via mTOR/P70S6K activation *in vitro*. However, the involvement of mTOR/P70S6K pathway in zinc-induced oxidative stress, tau degeneration, synaptic and cognitive impairment, has not been fully elucidated *in vivo*. Here we assessed in the effect of pathological concentration of zinc in SH-SY5Y cells by using biochemical assays and immunofluorescence staining. Rats (n = 18, male) were lateral ventricularly-injected with zinc and treated with rapamycin (intraperitoneal injection) for one week and assessed using Morris water maze. Evaluation of the oxidative stress, tau phorsphylation and synaptic impairment were performed using the hippocampus tissue of the rats by biochemical assays and immunofluorescence staining. Results from Morris water maze showed that the capacity of spatial memory is impaired in zinc-treated rats. Zinc sulfate significantly increased the levels of P-mTOR Ser2448, P-p70S6K Thr389, and P-tau Ser356, and decreased levels of Nrf2 and HO□1 in SH-SY5Y cells and in zinc-treated rats compared with control groups. Increased expressions of reactive oxygen species were observed in zinc sulfate-induced SH-SY5Y cells as well as in the hippocampus of zinc-injected rats. Rapamycin, an inhibitor of mTOR, rescued the zinc-induced increases in mTOR/p70S6K activations, tau phosphorylation and oxidative stress, as well as Nrf2/HO□1 inactivation, cognitive impairment and synaptic impairment reduced the expression of synapse-related proteins in zinc-injected rats. In conclusion, our findings imply that rapamycin prevents zinc-induced cognitive impairment and protects neurons from tau pathology, oxidative stress and synaptic impairment, by decreasing mTOR/p70S6K hyperactivity and increasing Nrf2/HO □ 1 activity.

## 1 Introduction

Alzheimer’s disease (AD) is pathologically featured by abnormal accumulation of amyloid-beta (Aβ) plaque, neurofibrillary tangles (NFTs), neuroinflammation, oxidative stress, synaptic impairment and neurodegeneration (1, 2). The pathological changes lead to cognitive decline and impairment in patients and in animal models (3, 4). The microtubule-associated protein tau is abnormally hyperphosphorylated and mainly aggregated into paired helical filaments (PHF) in brain from patients with AD (5, 6). Tau hyperphosphorylation is mediated by protein kinases or phosphatases, which involves in AD neurofibrillary degeneration (7, 8). The mammalian targets of rapamycin (mTOR) and ribosomal S6 protein kinase (p70S6K) are serine/threonine kinases, which play key roles in the regulation of protein synthesis and degradation, to age-dependent cognitive decline and pathogenesis of AD (9-11). Accumulating evidence demonstrated abnormal mTOR signaling in the brain affects several pathways in AD which associated with metabolism, insulin signaling, protein aggregation, mitochondrial function and oxidative stress (12). Increased expressions of mTOR and P70S6K being colocalized with NFT and mediated tau phosphorylation (9-11, 13, 14). Rapamycin, a well-known inhibitor of mTOR, plays an important role for autophagy and insulin signaling (15, 16) and regulating of tau phosphorylation (13, 17, 18). However, the upstream or downstream effectors that controlled by mTOR which contributed to the changes in neuronal functions and cognitive decline have not been fully elucidated.

Metal dysregulation, particularly iron, copper and zinc, is implicated in the development of AD at early stage (19-25). Higher level of cerebral zinc is observed in the post-mortem brain tissue from patients with AD (reaching 200–300 μM) compared to that in healthy controls (26, 27). Zinc is the second abundant essential trace metals in the brain, critical for the maintaining the brain homeostasis (19, 27, 28). Under pathological conditions, the excessive zinc released from the synaptic vesicle activation promotes tau hyperphosphorylation (19, 26, 29, 30) in cells (13, 31) and liquid-liquid phase separation of tau protein (32). In addition, previous studies have demonstrated that zinc firmly bind to Aβ and was detected inside Aβ plaque (33-36). In AD brain, presynaptic neurons release excessive zinc, which causes oxidase activation in neurons, and exacerbates the pathological development, leading to neuronal death (29, 30, 37, 38).

Oxidative stress is an early event in AD and play an important role in AD pathogenesis (39). Elevated level of reactive oxygen species (ROS) was detected in post-mortem brain tissues from AD patients and animal models of AD (40-42). The activation of nuclear factor erythroid 2-related factor 2 (Nrf2)/heme oxygenase-1 (HO-1) pathway inhibits the progression of inflammation, reduces ROS production, and thus has been a potential therapeutic target for AD (43-46). There is a vicious cycle formed by excessive zinc, tau and oxidative stress: elevated levels of zinc raises the production of ROS in mitochondria. The oxidative stress increases zinc concentration and tau hyperphosphorylation. In addition, excessive zinc and hyperphosphorylated tau cause oxidative stress and neurotoxicity. Hyperphosphorylated tau damages microtubule function and develops oxidative stress (47, 48); An increased oxidative stress has been indicated to cause tau hyperphosphorylation (49, 50), and aggravate neuronal death (51). Oxidative stress has been previously been shown as the underlying mechanism for the activation of mTOR in AD (52). However the underlying mechanism of how excessive zinc links to tau degeneration remains unclear. In the current study, we hypothesized that pathological concentration of zinc could disturb the rapamycin-dependent mTOR/P70S6K and Nrf2/HO□1 pathways, leading to detrimental effects in oxidative stress, tau hyperphosphorylation, synaptic and cognitive impairment. To this end, we assessed the effect of rapamycin treatment on the zinc sulfate (300 μM) treated SH-SY5Y cells and lateral ventrally-injected rats.

## 2, Materials and Methods

### 2.1 Materials and antibodies

Zinc sulfate, rapamycin, trisaminomethane (Tris), radioimmunoprecipitation assay (RIPA), sodium dodecyl sulphate (SDS) buffer and protease inhibitor cocktail were obtained from Sigma Aldrich Co. (St, Louis, MO, USA). A Bradford kit was purchased from Bio-rad (California, USA). The primary antibodies employed in the present study, please refer to **Supplementary Table 1**.

### 2.2 Cell culture and treatment

The cell culture is prepared as described previously (13, 53). Human SH-SY5Y neuroblastoma cells were grown to 70-80 % confluence in 75 cm^2^ plastic culture flasks (Corning, China) in a mixture of 5 % CO2 and 95 % air at 37 °C, employing Dulbecco’s modified Eagle’s medium (DMEM)/F12 medium (1:1) supplemented with 10 % fetal bovine serum (FBS), 100 units/mL penicillin, and 100 mg/mL streptomycin. Prior to treating the SH-SY5Y cells with 300 μM zinc sulfate, the cultures were kept in free serum media. 300 μM zinc sulfate is chosen based on the results and protocol established from our previous study (54). The SH-SY5Y neuroblastoma cells were pretreated with 20 ng/mL rapamycin for 1 h, then incubated with 300 μM zinc sulfate for 4 h.

### 2.3 Animals

Eighteen Sprague–Dawley (SD) rats were included in this study (male, weight 250-300 g, 12 months-of-age, Guizhou experimental Animal Center in China). All rats were housed in ventilated cages under in a climate-controlled room (temperature: 22 ± 2 °C, humidity: 50 ± 5 %, 12 h light– dark cycle with lights on at 08:00). Food (Safe, sterilized) and water (softened, sterilized) were provided ad libitum. Poplar wood shavings were placed in cages as environmental enrichments. All experimental protocols were approved by the Guiyang regional Animal Care center and Ethics Committee.

### 2.4 Surgery and treatment

Timeline of the surgery, treatment and behavior testing are shown in **Supplementary Figure 1**. All rats were randomly assigned into three groups (control group, zinc group, and zinc+rapamycin group, n = 6 each group). Rats were deeply anesthetized with an initial dose of 5 % isoflurane in oxygen/air mixture (1:4, 1 L/min) and were maintained at 1.5 % isoflurane in oxygen/air mixture (1:4, 0.6 L/min). Anesthetized rats were placed on a stereotaxic apparatus (RWD Life Science, Shenzhen, China) and the coordinates for injection were 0.8 mm posterior and 1.5 mm lateral and 3.6 mm ventral from the bregma. Zinc sulfate (25 mM, 2 μl) was injected slowly into the right lateral ventricle in rats from both Zn and Zn + Rapamycin group (**Figure 1A)**. Rats in control group underwent the same surgical procedures and were injected with phosphate-buffered saline (PBS, pH 7.4) of the same volume, respectively. Body temperature and respiratory rate of the rats were monitored during surgery. Body temperature of the animal was maintained at 36.5 ± 0.5 °C throughout the procedure using a warming pad. Lidocaine ointment was wiped locally to the scalp to reduce the pain. One day after the surgery, rats in the zinc+rapamycin group were administered with rapamycin (1.5 mg/kg body weight, intraperitoneal injection (i.p,), three times for one week). The rats in the control and Zn group were injected with 0.9 % citrate buffer of the same volume (i.p.). Behavioral tests were performed subsequently after rapamycin treatment period.

**Figure 1.**
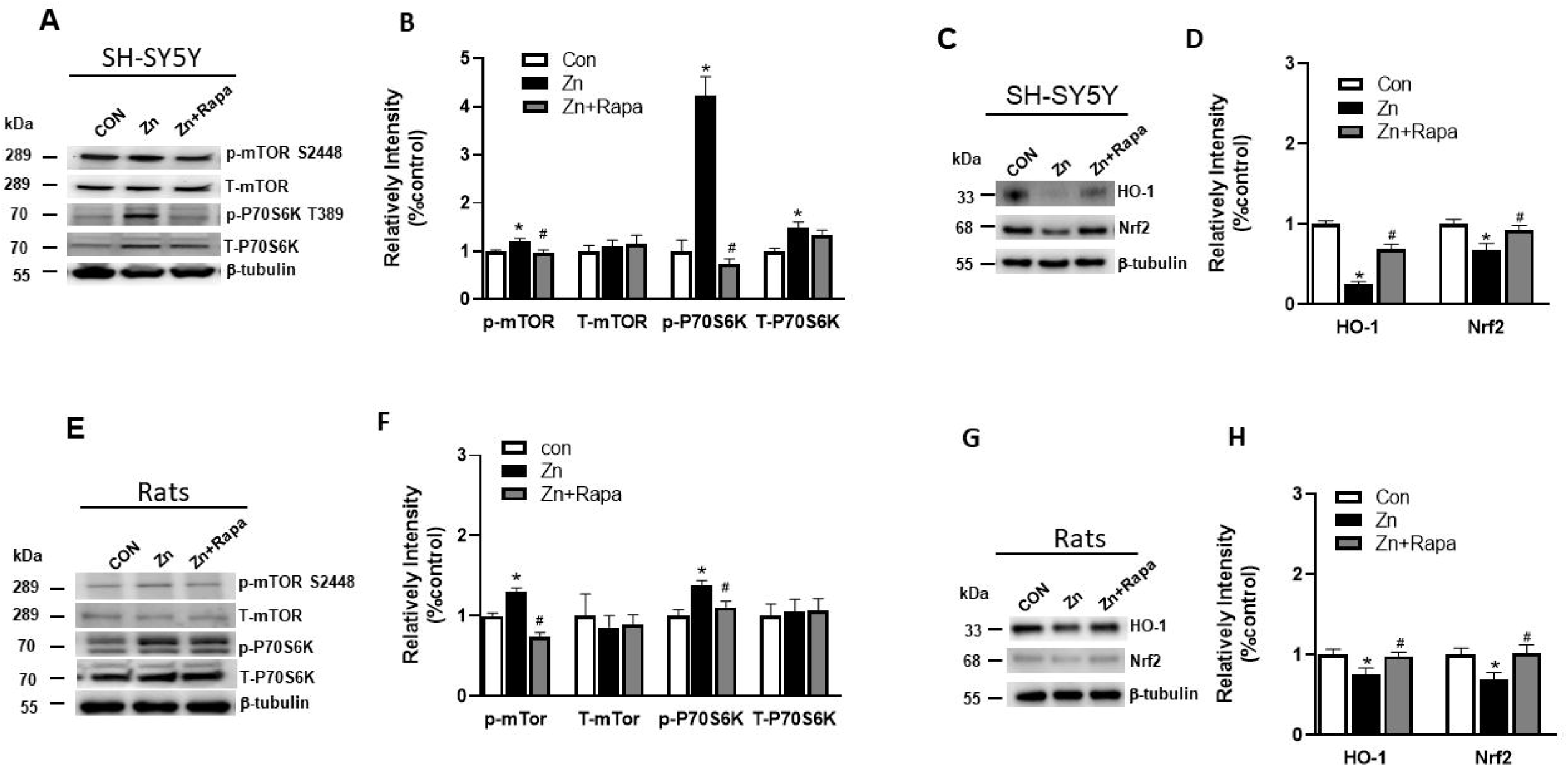
Effects of rapamycin on the expressions of mTOR/P70S6K and Nrf2/HO-1 signaling pathways in zinc treated SH-SY5Y cells and rat. (A, B) Representative blots and quantification of expression levels of p-mTOR S2448, T-mTOR, p-P70S6K T389, T-P70S6K in SH-SY5Y cells of control, zinc and zinc+rapamycin groups, n = 3 cell experiment per group. (C, D) Representative blots and quantification of expression levels of HO-1 and Nrf2 are detected in cells of control, zinc and zinc+rapamycin groups. n = 3 cell experiment per group; (E, F) Representative blots and quantification of expression levels of p-mTOR S2448, T-mTOR, p-nP70S6K T389, T-P70S6K in rats of control, zinc and zinc+rapamycin groups. n = 4 rats per group in the same membrane. (G, H) Representative blots and quantification of expression levels of HO-1 and Nrf2 in rats of control, zinc and zinc+rapamycin groups. n = 4 rats per group in the same membrane. Quantifications of the blots were normalized to β-tubulin. *p < 0.05 *vs*. control group; #p < 0.05 *vs*. zinc+rapamycin treatment.

### 2.5 mBehavioral Testing

Morris water maze (MWM) was used to assess the hippocampal spatial learning function of the rats (55). The circular pool (160 cm diameter and 50 cm height) was filled to a depth of 30 cm with water (25 ± 1 °C) in this study. Visual cues were positioned above water level and extra maze cues were blocked with a dark curtain. During Morris water maze training, all rats were subjected to 4 training trials daily for six consecutive days. In each trial, rats were trained to find a hidden platform (20 cm diameter) which submerged 1 cm under the water surface for 60 s. Afterwards the rats were kept to stay on the platform for 20 s. If these rats could not seek the platform within 60 s, they were guided to the platform within 60 s and kept to stay on the platform for 20 s afterwards. On day 7, a spatial probe trial was executed, where the platform was removed. The escape latency, the total of time spent in the target quadrant, the number of crossing platform, and swimming speed was monitored by video tracking software (ANY-maze, USA).

After the behavioral tests, all rats were then sacrificed under deep anesthesia with pentobarbital sodium (50 mg/kg body weight), and transcardiacly perfused with PBS (pH 7.4). Brains were removed from the skull afterwards. The left hemisphere brain tissue was saved for western blot and stored at -80 °C. The right hemisphere rat brain tissue was fixed in 4 % paraformaldehyde in 1 × PBS (pH 7.4) for 24 h and saved in 1 × PBS (pH 7.4) at 4 °C (56), For immunofluorescent staining, the fixed right brain hemisphere tissues were dehydrated using vacuum infiltration processor (Leica ASP200S, Germany), and embedded in paraffin using the Arcadia H heated embedding workstation (Leica, Germany).

### 2.6 Protein extraction and western blotting

The SH-SY5Y cells (n = 3 cell samples in each group) and the hippocampus of rats (n = 6 in each group) were lysed in RIPA buffer with a 0.1 % protease inhibitors cocktail on ice. Protein concentration was measured by a Bradford kit (Bio-rad). The proteins were analyzed by Western blotting as described earlier (13). The lysates were separated on 7.5-15 % SDS-PAGE gel, and the bands transferred onto 0.22/0.45 μm polyvinylidene difluoride (PVDF) membranes. After blocking the membranes with 5 % milk, the membranes were incubated with primary antibodies (**Supplementary Table 1**) at 4 °C overnight. The PVDF membranes were washed, and then with secondary peroxidase coupled anti-mouse or anti-rabbit antibodies (1:5000) at room temperature for 1 h (13). Immunoreactive bands were visualized by Immobilon Western Horseradish peroxidase substrate luminol reagent (Millipore) using a ChemiDoc™ MP imaging system (Bio-rad, USA).

### 2.7 DCFH-DA staining

ROS generation was measured via 2’-7’dichlorofluorescin diacetate (DCFH-DA) staining for detecting the intracellular hydrogen peroxide and oxidative stress (Beyotime, China). Following treatment and washing with PBS, the SH-SY5Y cells were incubated with DCFH-DA probes at 37 °C for 30 min. The intracellular accumulation of fluorescent DCF was imaged using by confocal microscopy (Leica, SP8, Germany).

### 2.8 Immunofluorescent staining and confocal imaging

After treatment, the SH-SY5Y cells were plated on coverslips were rinsed with PBS, and then were fixed in 4 % paraformaldehyde for 30 min. Cells were permeabilized in 0.1 % Triton X-100 in Tris-buffered saline (TBS) for 10 min. The unspecific binding sites were blocked with blocking solution (5% bovine serum albumin, 0.1 % Triton X-100 in TBS) for 1 h. Cells were incubated with primary antibodies, anti-4-hydroxynonenal (4-HNE) and anti-8-hydroxy-2’-deoxyguanosine (8-OHdG), at 4 □ overnight. After washing with TBS, bound antibody was detected by incubation for 1 h with Alexa Fluor 546-IgGs or Alexa Fluor 488-IgGs (1:200 for both, Invitrogen, USA).

Coronal sections of the rat brains were cut at 6 μm using a microtome (Leica RM2245, Germany). Dewaxed and rehydrated hippocampal sections were blocked in TBST (TBS with Tween20) with 5 % bovine serum albumin for 1 h, then incubated with the primary antibody against 8-OHdG at 4 °C overnight. After washing, the sections were incubated with AlexaFluor488 anti-mouse IgGs (1:200, Invitrogen, USA) for 1 h. After washing with TBS, the sections and coverslips were mounted by vector anti-fading mounting medium (Vector Laboratories, Burlingame, CA, USA). The fluorescent intensity was imaged using Leica SP8 confocal microscopy at 40 × magnification. The mean fluorescent intensity was quantified using Image J 1.49V software (NIH, US).

### 2.9 Statistical analysis

Statistical analysis was executed using SPSS software (version 23.0) or GraphPad Prism 8.0 software. The results were presented as mean ± SEM. Morris water maze behavior data were assessed by two-way repeated measure ANOVA. The other parameters were executed using one-way ANOVA followed by LSD’s *post-hoc* test for multiple comparisons. Significance was set at p < 0.05.

## 3 Results

### 3.1 Downregulation of mTOR/P70S6K and upregulation of Nrf2/HO_D_1 pathways are involved in the protection by rapamycin against the toxic effects of zinc sulfate both in SH-SY5Y cells and in rats

First, we assess the potential involvement of mTOR/P70S6K pathway in the alterations by zinc sulfate treatment in SH-SY5Y cells and zinc sulfate-injected rats. Zinc sulfate significantly elevated the levels of phosphorylated mTOR (S2448) by around 20 %, and phosphorylated P70S6K (T389) by 300 % in SH-SY5Y cells p = 0.015 and p = 0.0001 respectively, zinc sulfate-treated compared to control group (**Figs. 1A, B**). Pre-treatment with rapamycin (20 ng/mL) abolished the effect of zinc sulfate on the levels of mTOR (S2448), and phosphorylated P70S6K (T389) in SH□SY5Y cells, p = 0.01 and p = 0.0001 respectively, zinc-treated compared to zinc+rapamycin treated group (**Figures 1A, B**). The total levels of mTOR remained unaltered in the presence of zinc as well as zinc+rapamycin pretreatment in cell (**Figures 1A, B)**. Additionally, we found that zinc sulfate treatment induced an increase of total P70S6K protein in the zinc sulfate-treated cells compared with control cells, p = 0.01 (**Figure 1A**). Rapamycin pretreatment (20 ng/mL) did not affect the increased levels of total P70S6K protein induced by zinc sulfate in cells.

Zinc sulfate (300 μM) significantly elevated the levels of phosphorylated mTOR (S2448), and phosphorylated P70S6K (T389) by 25 % and 30 % in the hippocampus of the zinc-injected rats compared to control, p = 0.001, p = 0.005 respectively (**Figures 1E, F**). Rapamycin treatment attenuated the effect of zinc sulfate on mTOR (S2448), and phosphorylated P70S6K (T389) in rats, p = 0.0001, p = 0.026 (zinc *vs*. zinc+rapamycin group) (**Figures 1E, F**). Both the levels of total mTOR and total P70S6K protein remained unaltered in the presence of zinc sulfate as well as zinc+ rapamycin pretreatment in all three groups of rats (**Figures 1E, F)**.

Next, we assess the potential involvement of Nrf2 and HO□1 pathway in the zinc induced alterations using SH-SY5Y cells and zinc-induced rats (**Figures 1C-D, G-H**). Zinc sulfate (300 μM) significantly reduced the levels of Nrf2 by approximately 70 %, and HO□1 by 30 % in SH□SY5Y cells p = 0.0003, and p = 0.016 (**Figures 1C, D**). Pre-treatment with rapamycin (20 ng/mL) attenuated the effect of zinc sulfate on levels of Nrf2, and HO□1, p = 0.001 and p = 0.043 in SH[SY5Y cells, zinc-treated compared to zinc+rapamycin treated group (**Figures 1C, D**).

In the hippocampus of zinc-injected rats, zinc sulfate (300 μM) significantly reduced the levels of Nrf2 and HO □ 1 by approximately 20% and 30 %, p = 0.031 and p = 0.04, zinc injected *vs*. zinc+rapamycin group (**Figures 1G, H**). Rapamycin treatment abolished the effect of zinc sulfate on levels of Nrf2, and HO □ 1, p = 0.04 and p = 0.04 in rats, zinc injected *vs*. zinc+rapamycin group (**Figures 1G, H**). Hence the neuroprotective effects of rapamycin were linked to the inactivation of the mTOR/P70S6K and activation of the Nrf2/HO □ 1 signaling pathways as defensive responses to oxidative stress.

### 3.2 Rapamycin ameliorates tau hyperphosphorylation in zinc-induced SH-SY5Y cells and rats

Next we assess the potential rapamycin protection against zinc induced tau hyperphosphorylation using SH-SY5Y cells and zinc-injected rats. We found that zinc treatment led to increased level of hyperphosphorylated tau at Ser356 (by 100 %) in zinc-treated SH-SY5Y cells compared to control, p = 0.005. Rapamycin treatment completely restored the level of phosphorylated tau S356 in SH-SY5Y cells, p = 0.003 zinc-treated compared to zinc+rapamycin treated group. The total tau levels showed no change in the presence of zinc or zinc+rapamycin compared to control group in cells (**Figures 2A, B**).

**Figure 2.**
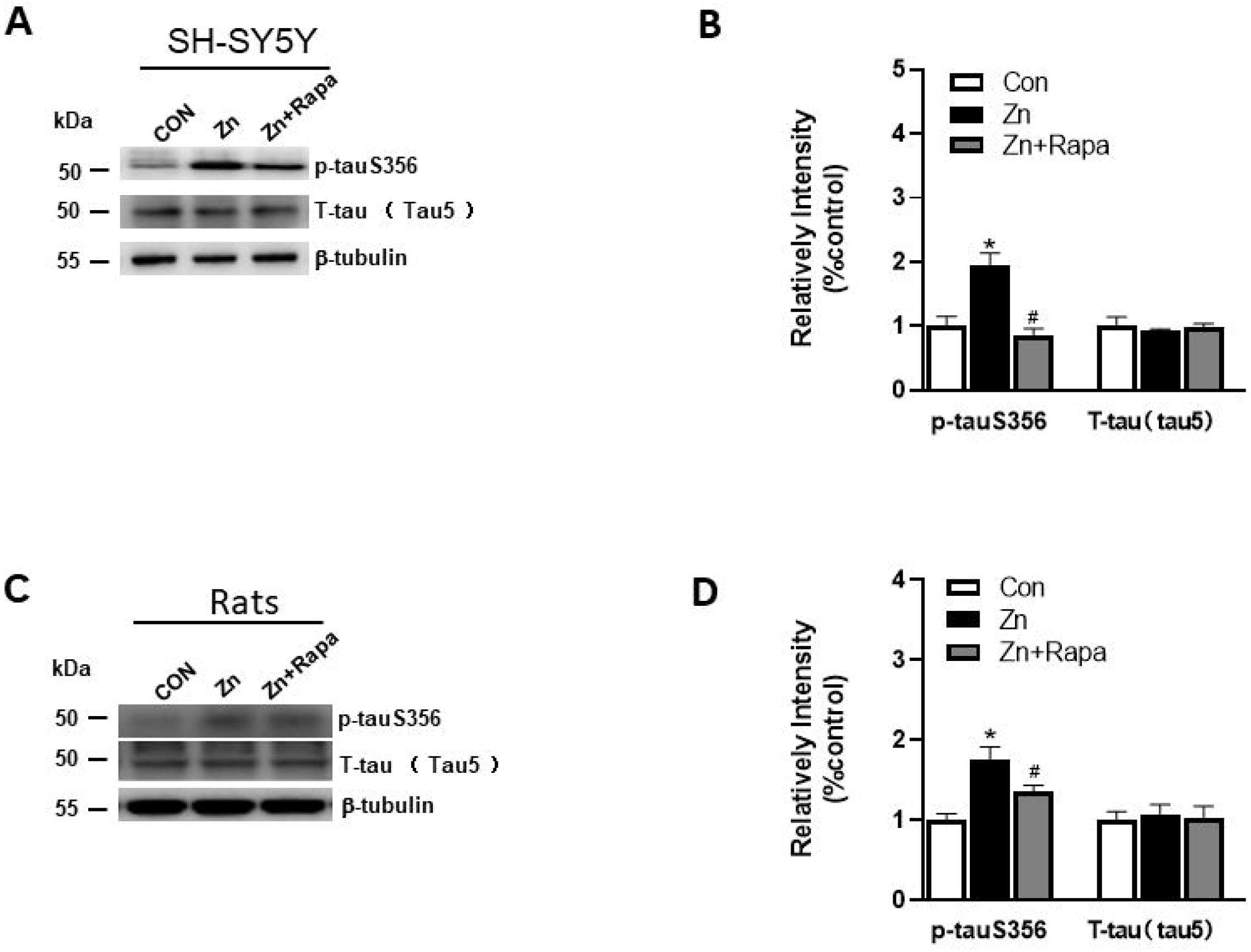
Rapamycin decreases the expression of p-tau S356 in SH-SY5Y cells. (A, B) Representative blots and quantification of expression levels of p-Tau S356 and T-tau (tau5) proteins in SH-SY5Y cells of control, zinc and zinc+rapamycin groups. n = 3 cell experiment per group; (C, D) Representative blots and quantification of expression levels of p-Tau S356, T-tau (tau5) protein in rats of control, zinc and zinc+rapamycin groups. n = 4 rats per group in the same membrane. *p < 0.05 *vs*. control group; #p < 0.05 *vs*. rapamycin treatment. Quantifications of the blots were normalized to β-tubulin.

Moreover, zinc led to increased level of hyperphosphorylated tau at Ser356 in the hippocampus of zinc-injected rats by 80 %, p = 0.001 (**Figure 2C, D**). Rapamycin treatment decreased the level of phosphorylated tau S356 by 40 % in zinc-injected rats compared to zinc+rapamycin group, p = 0.031. The total tau levels showed no change in the presence of zinc or zinc+rapamycin in rats (**Figures 2C, D**).

### 3.3 Rapamycin attenuates oxidative stress damage in SH-SY5Y cells and in the hippocampus of zinc induced rats

Next we assessed the oxidative stress by immunofluorescence staining using DCFH-DA, 4□HNE (lipid peroxidation), and 8□OHdG (oxidation of DNA) in SH-SY5Y cells. Following exposure to zinc sulfate, the level of fluorescence intensity of DCFH-DA, 4□HNE and 8□OHdG elevated in the zinc-treated group compared to the control group in SH-SY5Y cells, p = 0.0001, p = 0.0001, p = 0.0001 respectively (**Figures 3A, B, 4A, B** and **5A, B**). Prior treatment with rapamycin neutralized the zinc-induced increase in fluorescence intensity of DCFH-DA, 4□HNE and 8□OHdG, zinc-treated group compared to zinc+rapamycin group in SH-SY5Y cells, p = 0.0004, p = 0.0003, p = 0.0004 respectively (**Figures 3A, B, 4A, B** and **5A, B**).

**Figure 3.**
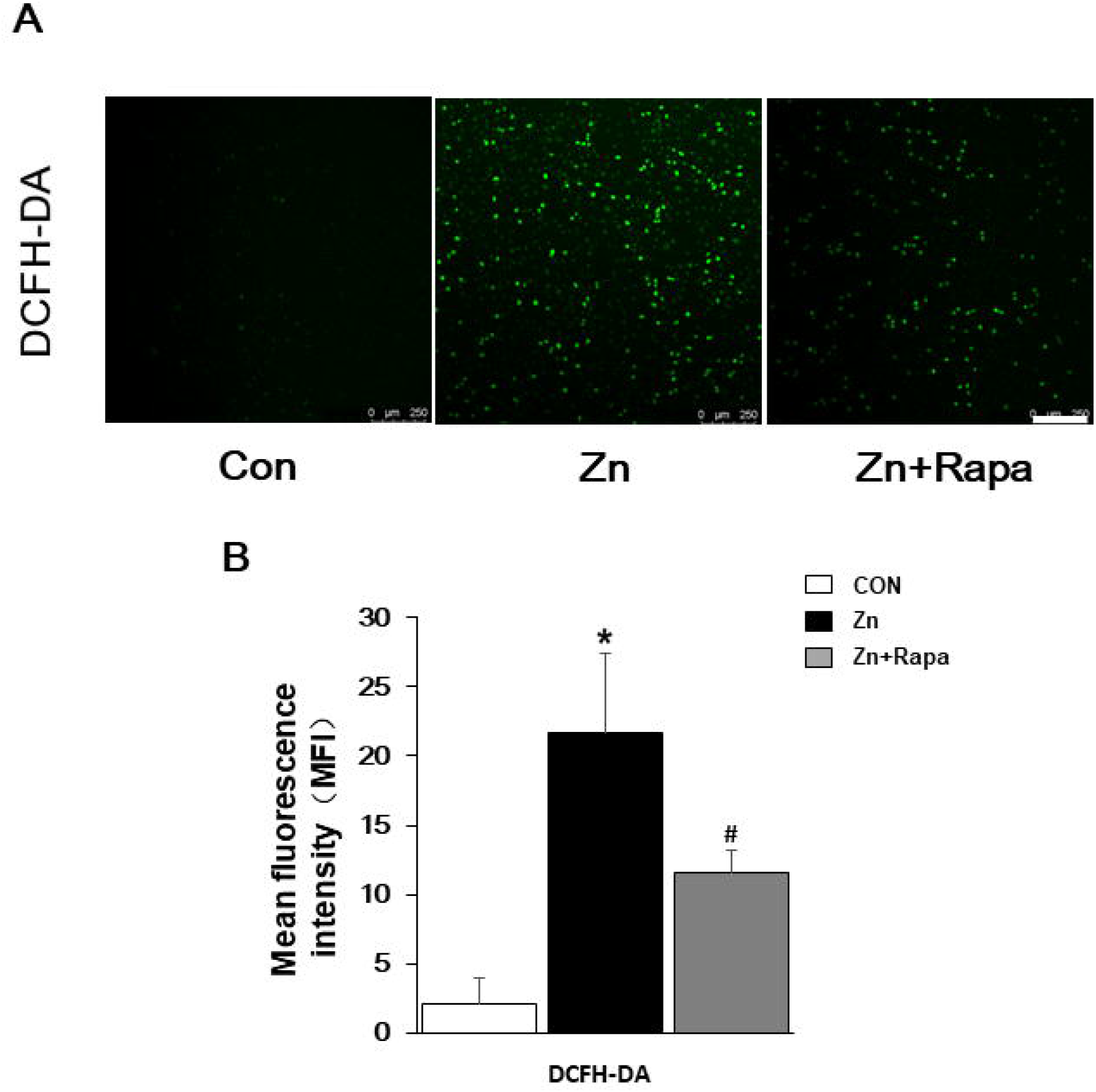
Rapamycin suppressed ROS generation caused by zinc in SH-SY5Y cells. (A, B) Representative fluorescent staining and quantification of the mean fluorescence intensity of ROS determined by DCFH-DA staining in SH-SY5Y cells of control, zinc and zinc+rapamycin groups. Scale bar = 250 μm.; n = 3 cell experiment per group,*p < 0.05 *vs*. control group; #p < 0.05 *vs*. zinc+rapamycin treatment.

To determine oxidative stress damage induced by zinc *in vivo* in rats, we measured the levels of 4-HNE by using western blot in the hippocampal brain tissue homogenates from rats. Zinc led to an increased level of 4-HNE in the hippocampus of zinc-injected rats, p = 0.005, compared to control group (**Figure 4C, D**). Rapamycin treatment decreased the level of 4-HNE in zinc-injected rats compared to zinc+rapamycin group, p = 0.04 (**Figures 4C, D**). We further measured the levels of 8-OHdG products by immunofluorescence staining using in the hippocampal brain tissue slices from rats. Zinc led to an increased level of 8-OHdG in the hippocampus of zinc-injected rats, p = 0.001 compared to control group (**Figure 5C, D**). Rapamycin treatment decreased the level of 4-HNE, and 8-OHdG by in zinc-injected rats compared to zinc+rapamycin group, %, p = 0.038 (**Figures 5C, D**).

**Figure 4.**
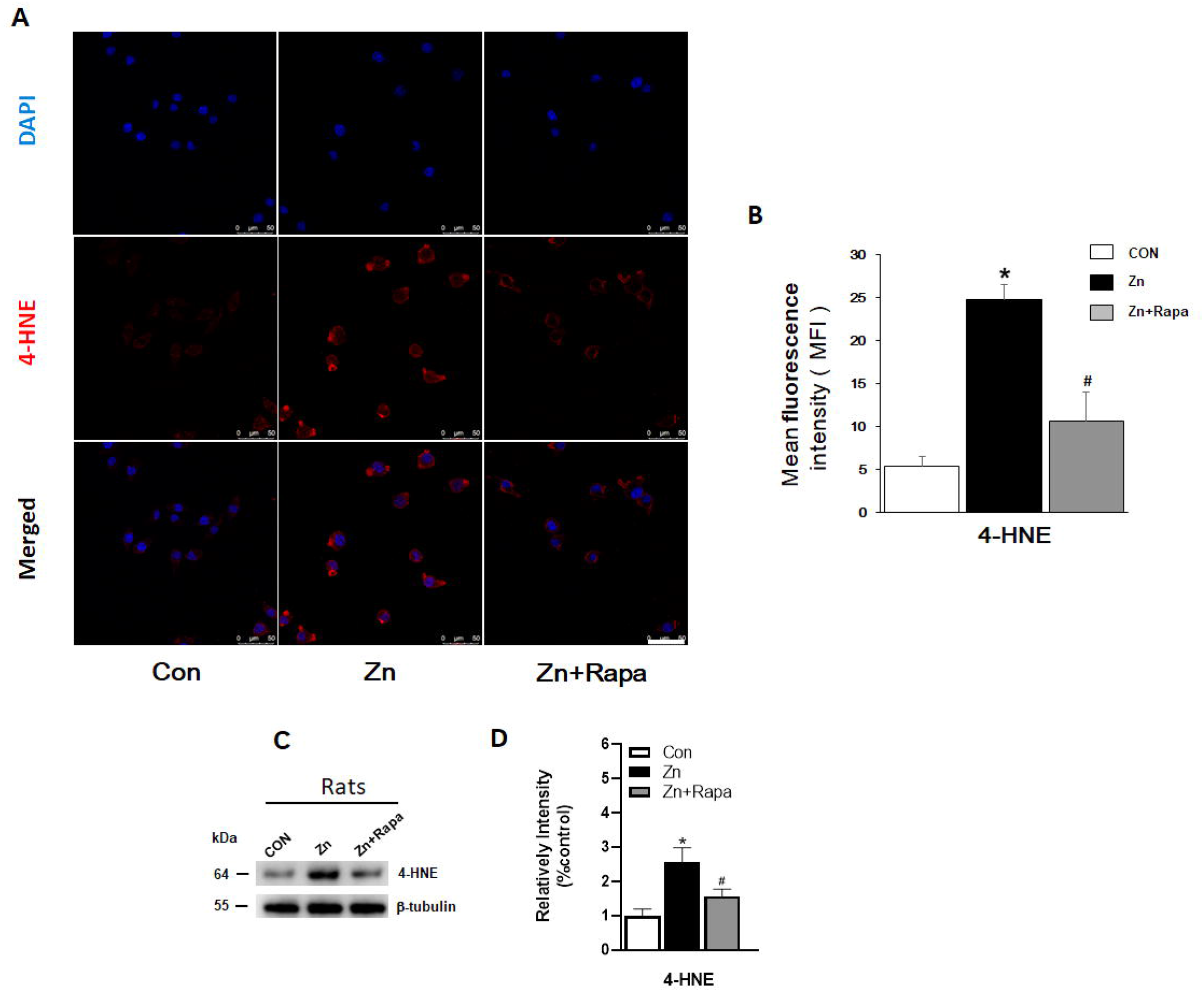
Rapamycin attenuated the increase in the expressions of 4-HNE caused by zinc both *in vitro* and *in vivo*. (A, B) Representative confocal images and quantification of the mean fluorescence intensity for immunofluorescence staining using anti-4-HNE antibody (red) in SH-SY5Y cells in control, zinc and zinc+rapamycin groups, the scale bar = 50 μm; Nuclei is counter-stained with DAPI (blue); (C, D) Representative blots and quantification of expression levels of 4-HNE in rats of control, zinc and zinc+rapamycin groups. n = 4 rats per group in the same membrane. *p < 0.05 *vs*. control group; #p < 0.05 *vs*. rapamycin treatment. Quantifications of the blots were normalized to β-tubulin.

**Figure 5.**
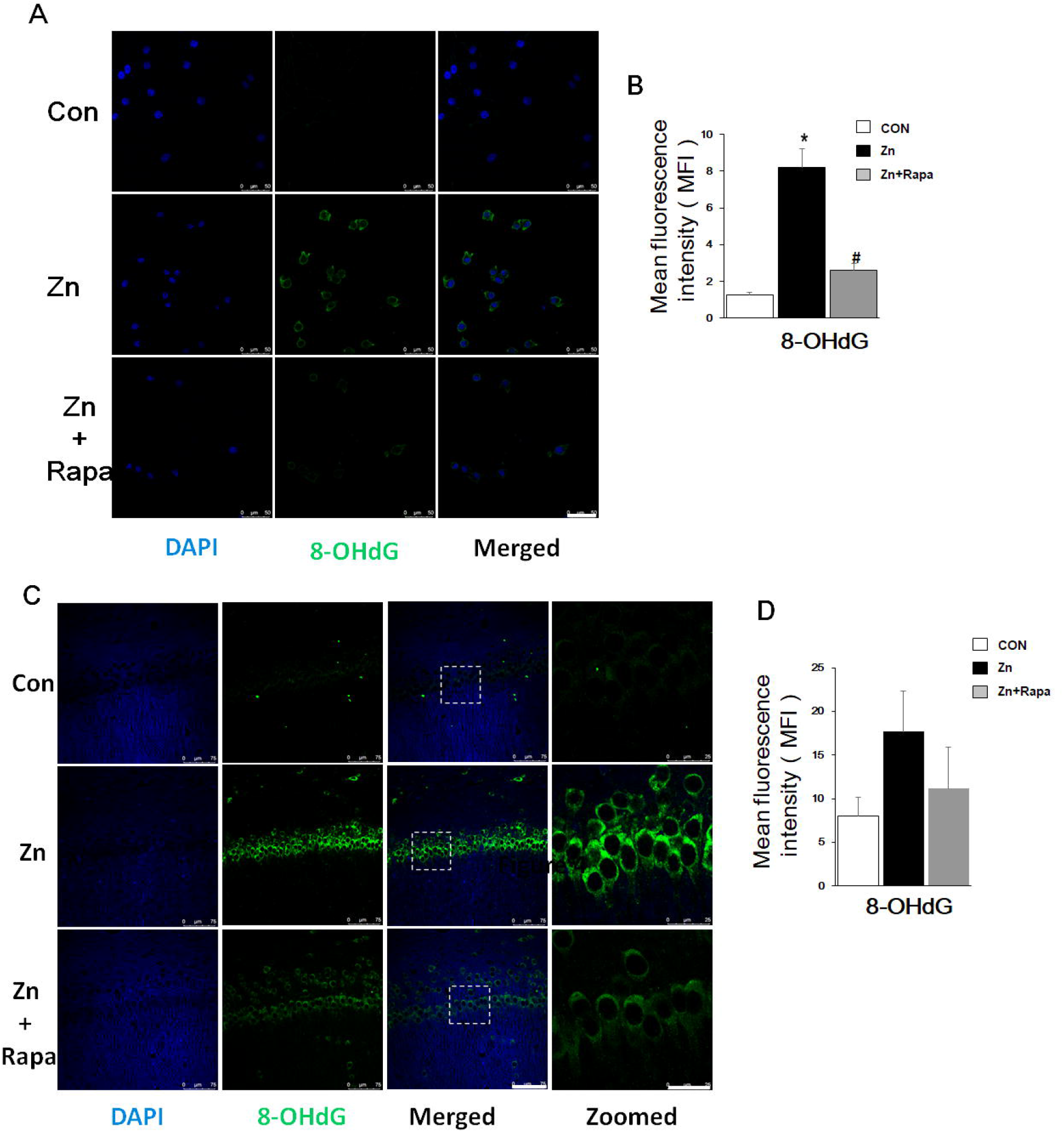
Rapamycin prevents the DNA oxidation caused by zinc in SH-SY5Y cells and rats. (A, B) Representative confocal images and quantification of the mean fluorescence intensity for immunofluorescence staining using anti-8-OHdG antibody (green) in SH-SY5Y cells in control, zinc and zinc+rapamycin groups. Scale bar = 50 μm; Nuclei is counter-stained with DAPI (blue); Quantification of the mean fluorescent intensity of 8-OHdG staining in the SH-SY5Y cells of control, zinc and zinc+rapamycin groups, n = 3 cell experiment per group; (C, D) Representative confocal images and quantification of the mean fluorescence intensity for immunofluorescence staining using anti-8-OHdG antibody (green) in the CA1 areas of the brain of the rats in control, zinc and zinc+rapamycin groups. Nuclei is counter-stained with DAPI (blue). Scale bar = 75 μm, and 25 μm (zoomed images); n = 6 rats per group. *p < 0.05 *vs*. control group; #p < 0.05 *vs*. zinc+rapamycin treatment.

### 3.5 Rapamycin rescued impaired learning and memory in zinc-induced rats

The spatial learning and memory function of the rats were assessed by using MWM test. The escape latency significantly increased in the zinc-injected rats compared to control on Day 5 and Day 6 (p = 0.031, p =0.024 respectively) (**Figure 6A**). On Day 7, the time spent in the target quadrant and the number of platform location crossings both decreased by approximately 50 % in zinc-induced rats compared to the control group (p = 0.027 and p = 0.039) (**Figures 6B, D**).

**Figure 6.**
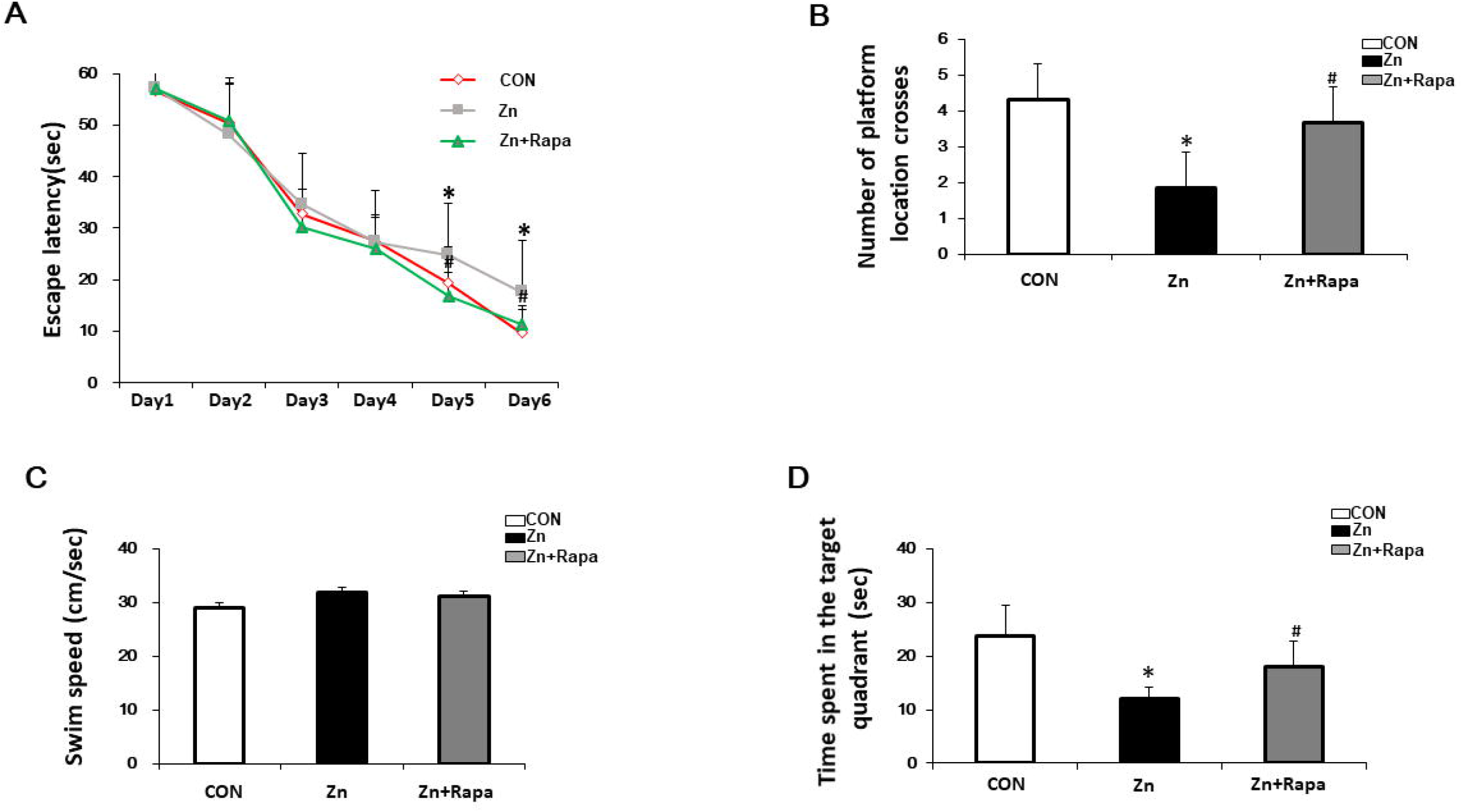
The effect of rapamycin on the performance of MWM in zinc-induced rats. (A) Escape latency of the rats in all three groups (control (red), zinc-injected (grey), zinc+rapamycin treatment (green)) in the MWM test of each training day. (B) Representative number of platform crossings on day 7. (C) Representative time spent in the target quadrant on day 7. (D) Swimming speed in MWM on day 7. n = 6 rats per group, *p < 0.05 *vs*. control group; #p < 0.05 *vs*. rapamycin treatment.

Treatment with rapamycin led to a reduced escape latency in zinc + rapamycin rats compared to the zinc-injected rats on Day 5 and Day 6 (p = 0.021, p = 0.033 respectively). On Day 7, the time spent in the target quadrant and the numbers of platform location crossings are increased compared to zinc-injected rats (p = 0.044, p = 0.044), (**Figures 6B, D**). To study if zinc could affect the motion ability of rats, the swimming speed of rats were recorded. No differences were observed among the three groups (**Figure 6C**), implying that treatment with rapamycin and zinc did not radically affect the motion ability of rats.

### 3.6 Rapamycin protected the synapses in zinc-induced SH-SY5Y cells and in rats

Next we assessed the potential effect of rapamycin on zinc-induced synaptic impairment using SH-SY5Y cells and zinc-induced rats. The expression levels of presynaptic proteins (SNAP 25 and synaptophysin) and postsynaptic protein (PSD 95) are indicator of the synaptic function. We found that zinc treatment led to reduced levels of SNAP 25, synaptophysin and PSD 95 in SH-SY5Y cells zinc treated compared to control, p = 0.039, p = 0.004 and p = 0.04 respectively. Rapamycin treatment reversed the reduction in the levels of synaptophysin and PSD 95 induced by zinc in SH-SY5Y cells p = 0.03, p = 0.0001, and p = 0.035 (zinc-treated group compared to zinc+rapamycin group)(**Figures 7A, B**). The levels of SNAP 25 recovered back to the control level, although were not significantly different from the zinc-treated cells.

**Figure 7.**
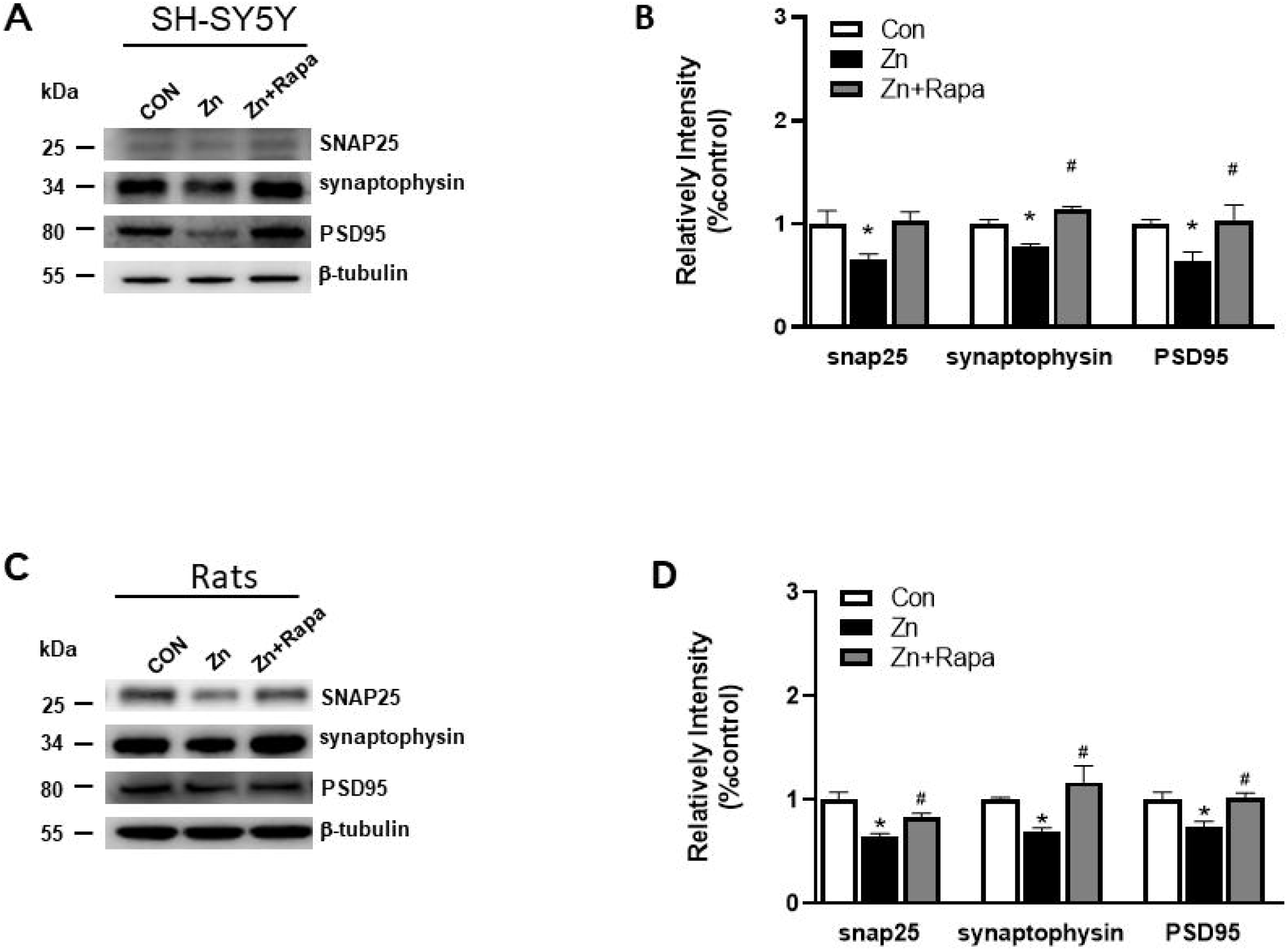
Rapamycin improved synaptic impairment caused by zinc both *in* SH-SY5Y cells and in *rats*. (A, B) Representative blots and quantification of expression levels of SNAP25, synaptophysin and PSD95 protein expressions in SH-SY5Y cells of control, zinc and zinc+rapamycin groups. n = 3 cell experiment per group; (C, D) Representative blots and quantification of expression levels of SNAP25, synaptophysin and PSD95 protein expressions in rats of control, zinc and zinc+rapamycin groups. n = 4 rats per group in the same membrane, *p < 0.05 *vs*. control group; #p < 0.05 *vs*. zinc+rapamycin treatment. Quantifications of the blots were normalized to β-tubulin.

Moreover, zinc injection led to reduced levels of SNAP 25, synaptophysin and PSD 95 in the hippocampus of zinc-injected rats, p = 0.001, p = 0.04, p = 0.013 respectively (**Figures 7C, D**). Rapamycin treatment decreased the level of SNAP 25, synaptophysin and PSD 95 in zinc-injected rats compared to zinc+rapamycin group, p = 0.038, p = 0.006, p = 0.009 respectively (**Figures 7C, D**).

## 4 Discussion

Our data reveal that zinc leads to tau hyperphosphorylation, oxidative stress and synaptic impairment involving mTOR/P70S6K activation and Nrf2/HO□1 inactivation. Rapamycin ameliorates the zinc-induced tau hyperphosphorylation, oxidative stress damage and synaptic impairment, as well as rescues spatial learning deficits via down-regulating mTOR/P70S6K activities and up-regulating Nrf2/HO □ 1 activities.

Zinc has been closely related to the cognitive function, play an important role under physiological condition. Elevated levels of zinc ion were found in AD brains, notably in the hippocampus, cortex, and amygdale, which are severely affected by NFTs (19, 57-59). Accumulation evidence have shown a tight relationship of zinc with tau degeneration and cognitive impairment in human patients (60). Dietary zinc supplementation treatment in 3xTg-AD mice has been shown increased BDNF levels and prevented cognitive deficits as well as mitochondrial dysfunction (61). Pathological level of zinc promotes tau tangle pathology in the brain of hAPP/htau (62), and tau mouse models (63). Targeting metals abnormal accumulation has been shown to rescue the pathology and phenotype in transgenic mouse models of tauopathy (64).

In the present study, we found that excessive zinc could induce dependent-mTOR(S2448)-P70S6K(T389) phosphorylation and tau hyperphosphorylation both in cultured neuroblastoma SH-SY5Y cells. Rapamycin suppress mTOR(S2448)/P70S6K(T389) phosphorylation and ameliorate tau pathology. We found that lateral ventricular injection of zinc sulfate could induce a persistent mTOR(S2448)-P70S6K(T389) phosphorylation in the rats’ hippocampus, and that rapamycin reversed both mTOR(S2448) and P70S6K(T389) phosphorylation, and reduced the level of tau hyperphosphorylation. These results implied that zinc played a crucial role in tau pathology via regulating mTOR/P70S6K pathway and rapamycin exerted a beneficial effect on tau pathology.

mTOR or P70S6K is one of the most important serine/threonine kinase in eukaryotic cells, which plays a prominent role in the regulation of protein synthesis, phosphorylation, and autophagy (11, 65-69). We have previously showed increased expressions of p-mTOR(S2448) and p-P70S6K(T389) in post-mortem AD brains, which associated with accumulation of hyperphosphorylated tau in AD (13, 14, 70). This suggests that phosphorylation of mTOR at S2448 and P70S6K at T389 is a vital target for disease intervention. Zinc has been indicated in the mechanisms of mTOR/P70S6K activation in AD. We have previously shown that zinc treatment (300 μM) promoted tau phosphorylation *in vitro* in cell cultures (13). Other study demonstrated that synaptic zinc promoted tau hyperphosphorylation (30), and accelerated the fibrillization of mutant ΔK280 of full-length human tau, inducing apoptosis and toxicity in SH-SY5Y cells via bridging Cys-291 and Cys-322 (71). In addition, zinc binds to protein phosphatase 2A and induces its inactivation and tau hyperphosphorylation through Src dependent PP2A (tyrosine 307) phosphorylation (72). In ApoE4 transgenic mice, the tau hyperphosphorylation is associated with activation of extracellular signal-regulated kinase that modulated by zinc (73).

Oxidative stress has been recognized as a causative factor in various neurodegenerative diseases (41, 42, 44, 45). Excessive zinc triggers oxidase activation, and further generates oxidative products in neurons (19, 27). Pathological tau damages mitochondrial function resulting in increased ROS products and causing oxidative stress (26, 27, 74, 75). Increased productions of ROS causes lipid peroxidation and DNA damage, and in turn, could affect the hyperphosphorylation of tau, leading to a vicious cycle (42, 49, 75-77). Here we investigate the mechanism of the zinc-mediated tau pathology in AD and whether excessive zinc could exacerbate the vicious cycle through mTOR/P70S6K pathway. The levels of oxidative products were assessed by immunostaining or western blot in the present study. We found an increasing level of ROS accumulation in zinc-treated SH-SY5Y cells, a marked increased expressions of lipid peroxidation product (4-HNE) and nucleic peroxidation product (8-OHdG) in both zinc-induced SH-SY5Y cells and zinc-injected rats. Employing rapamycin pre-administration, the level of ROS, 4-HNE and 8-OHdG was decreased in zinc-treated SH-SY5Y cells and zinc-injected rats.

Nrf2 is a transcription factor that negatively regulates the level of ROS to protect against oxidative stress damage. Ramsey et al. reported a significant decline in the level of Nrf2 in the brain from patients with AD (78). Several natural compounds havee been shown reduced oxidative stress in AD models through the Nrf2/HO-1 pathway (79, 80). We found rapamycin reverted the reduced level of Nrf2 /HO-1 in zinc-induced SH-SY5Y cells and rats, in line with previous observation (81). Rapamycin also rescued oxidative stress caused by tau hyperphosphorylation associated with inactive mTOR/P70S6K signaling pathway and active Nrf2/HO-1 signaling pathway.

The altered mTOR or P70S6K has been directly correlated to learning and memory in animal models (15, 82-84). Caccamo and Oddo et al. found that inhibition of mTOR by rapamycin could improve learning and memory, and reduce Aβ and tau pathology in 3×Tg-AD mice (85, 86). Chronic treatment with rapamycin enhances learning and memory in young adult mice, and improves age-related cognitive decline in older mice, possibly by activating major monoamine pathways in brain (87). In the present study, we show that an increased mTOR/P70S6K signaling in hippocampus of rat. Rapamycin treatment rescued the abnormal mTOR/P70S6K signaling to and improved the spatial learning in rats. Moreover we showed that rapamycin treatment reversed the decreased expression levels of synaptic proteins SNAP 25, synaptophysin and PSD 95 both in cell culture and in rats. Accumulating evidence have shown that **r**apamycin **improved the** cognitive decline in mouse model of Down syndrome **(15, 88, 89), amyloidosis** (J20) **(90-93), tauopathy (17, 94, 95)**. Despite its compelling preclinical record, no clinical trials have tested rapamycin in patients with **AD (96, 97)**.

There are several limitations in the study. Firstly, only MWM test was used to assess the spatial learning function of the rats. Further study using a panel of behaviour tests will provide comprehensive insights into the effect of rapamycin on zinc induced cognitive impairment. Secondly, we focused on the involvement of mTOR/p70S6K and Nrf2/HO-1 pathway in the current study, while many other pathways have been implicated in the effect elicited by zinc and rapamycin. Moreover, an acute effect of zinc and rapamycin treatment was investigated in the current study. The effect of chronic or environmental zinc exposure and treatment using rapamycin via oral intake remains to be investigated. The advance in non-invasive imaging has enabled detection of metal accumulation in vivo by using magnetic resonance imaging (98). Longitudinal study using *in vivo* imaging of the treatment effect on tau, neuroinflammation, ROS in animal model will provide further provide systematic insights (25, 54, 99-104).

## 5 Conclusions

In conclusion, zinc treatment induces mTOR/p70S6K activation and Nrf2/HO-1 inactivation, tau hyperphosphorylation, oxidative stress damage in SH-SY5Y cells and spatial learning impairment in rats. Rapamycin attenuate mTOR/p70S6k and increase Nrf2/HO-1 activity, attenuates tau pathology, oxidative stress and cognitive deficits induced by zinc in a rat model. Rapamycin might be a viable treatment for zinc related neuronal and synaptic damage.

## 6 Conflict of Interest

The authors declare that there is no conflict of interests regarding the publication of this paper.

## 7 Author Contributions

ZT contributed to the conception and study design. CCL, YTD and QC performed the experiments. HL and SBS contributed to data collection and data analysis. RN, ZT interpreted the data. CCL, YTD, RN, and ZT wrote the manuscript. All authors approved the manuscript before submission.

## 8 Funding

This work was supported by the Chinese National Natural Science Foundation (81560241), China Postdoctoral Science Foundation (2020M683659XB), and the Foundation for Science and Technology projects in Guizhou ([2020]1Y354), Scientific Research Project of Guizhou University of Traditional Chinese Medicine ([2019]48).

## Notes

### Competing Interest Statement

The authors have declared no competing interest.

