## Supplementary table 1 for "Rapamycin attenuated zinc-induced tau phosphorylation and oxidative stress in animal model: Involvement of dual mTOR/p70S6K and Nrf2/HO-1 pathways"

**Supplementary Table 1. Antibodies used in this study.**

| **Antibody** | **Host** | **Specificity** | **Phosphoepitopes** | **WB**  **Dilution** | | **IF Dilution** | **Sources** |
| --- | --- | --- | --- | --- | --- | --- | --- |
| Anti-Tau S356 | r | p-Tau | S356 | 1:1000 | | - | Invitrogen |
| Anti-Tau5 | m | T Tau5 |  | 1:500 | |  | Santa cruz |
| Anti-mTor (7C10) | r | T mTor | - | 1:1000 | | - | Cell Signaling |
| Anti-p-mTor | r | P, active mTor | S2448 | 1:1000 | | - | Cell Signaling |
| Anti-p-P70S6K (108D2) | r | P-P70S6K | T389 | 1:1000 | | - | Cell Signaling |
| Anti -P70S6K | r | T P70S6K |  | 1:1000 | | - | Cell Signaling |
| SNAP25 | r | SNAP25 | - | 1:1000 |  | | Abcam |
| synaptophysin | r | synaptophysin | - | 1:1000 |  | | Abcam |
| PSD95 | r | PSD95 | - | 1:1000 |  | | Abcam |
| 4-HNE | r | 4-HNE | - | 1:3000 | 1:100 | | Abcam |
| 8-OHdG | m | 8-OHdG | - | - | 1:200 | | Abcam |
| HO-1 | r | HO-1 | - | 1:1000 |  | | Abcam |
| Nrf2 | m | Nrf2 | - | 1:1000 |  | | Abcam |

T: total; r: rabbit; m: mouse; p: phosphorylated; IF: immunofluorescence; WB: western blot;
