## Supplementary figure 1 for "Rapamycin attenuated zinc-induced tau phosphorylation and oxidative stress in animal model: Involvement of dual mTOR/p70S6K and Nrf2/HO-1 pathways"

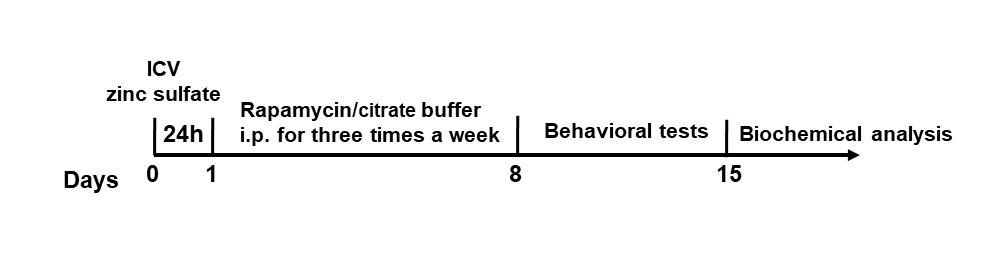

**Supplementary figure 1 Timeline of surgery, treatment and behavioral test in rats.** ICV: Intracerebroventricular
